# Decoding muscle-resident Schwann cell dynamics during neuromuscular junction remodeling

**DOI:** 10.1101/2023.10.06.561193

**Authors:** Steve D. Guzman, Ahmad Abu-Mahfouz, Carol S. Davis, Lloyd P. Ruiz, Peter C.D. Macpherson, Susan V. Brooks

## Abstract

This investigation leverages single-cell RNA sequencing (scRNA-Seq) to delineate the contributions of muscle-resident Schwann cells to neuromuscular junction (NMJ) remodeling by comparing a model of stable innervation with models of reinnervation following partial or complete denervation. The study discovered multiple distinct Schwann cell subtypes, including a novel terminal Schwann cell (tSC) subtype integral to the denervation-reinnervation cycle, identified by a transcriptomic signature indicative of cell migration and polarization. The data also characterizes three myelin Schwann cell subtypes, which are distinguished based on enrichment of genes associated with myelin production, mesenchymal differentiation or collagen synthesis. Importantly, SPP1 signaling emerges as a pivotal regulator of NMJ dynamics, promoting Schwann cell proliferation and muscle reinnervation across nerve injury models. These findings advance our understanding of NMJ maintenance and regeneration and underscore the therapeutic potential of targeting specific molecular pathways to treat neuromuscular and neurodegenerative disorders.

## Introduction

The neuromuscular junction (NMJ) is the terminal synapse of the motor system, orchestrating muscle contraction and, by extension, voluntary movement. Effective neurotransmission at the NMJ is paramount for neuromuscular communication, with any degenerative alterations at the synapse having the potential to impair muscle contractile function and overall mobility. The major cellular components of the NMJ are the pre-synaptic motor neuron (MN), the post-synaptic muscle fiber, and supporting glia. The primary glial cell type at the NMJ is a specialized non-myelinating Schwann cell population referred to as terminal Schwann cells (tSCs). tSCs regulate synaptic activity (*1*), and tSC proliferation is critical for effective muscle reinnervation (*2*, *3*). Despite the importance of Schwann cells (SC) and SC proliferation in promoting muscle fiber reinnervation, the cell biology and physiology of muscle resident SCs and their response to cues that induce reinnervation remain poorly understood.

To comprehensively assess the role of muscle-resident Schwann cells in NMJ remodeling and muscle fiber reinnervation, we conducted single-cell RNA sequencing (scRNA-Seq) on cells isolated from muscles during recovery from complete denervation post-surgical nerve crush injury and from 2-month-old *Sod1*^-/-^ mice undergoing spontaneous denervation and reinnervation. The *Sod1*^-/-^ mouse model, which lacks the enzyme superoxide dismutase 1 (Sod1), displays widespread and progressive NMJ degeneration and denervation and is thus a widely used model for testing hypotheses focused on the regulators of NMJ structure and function (*4–6*). A recent revelation from this mouse model is a distinct timeframe early in life during which known markers of muscle fiber denervation (*7–9*), acetylcholine receptor alpha (*Chrna*) mRNA levels and mitochondrial reactive oxygen species (mtROS) production, are dramatically but transiently elevated (*10*). A pattern of increases around 2 months of age and return to baseline by 3 months in *Sod1^-/-^* mice for both Chrna mRNA and mtROS suggests a robust denervation event that is followed by successful reinnervation and thus, identifies a ’regenerative window’ in which NMJ repair processes following spontaneous partial denervation can be examined alongside the full denervation associated with surgical nerve crush injury.

We first rigorously confirmed widespread denervation with a successful regeneration response between 1 and 3 months of age in *Sod1*^-/-^ mice supporting the use of young *Sod1*^-/-^ mice as a novel genetic model of spontaneous recoverable partial denervation. Our scRNA-Seq data revealed several Schwann cell subtypes in muscle, including a newly identified tSC subtype is enriched during the denervation-reinnervation response. This subtype is characterized by a unique transcriptomic signature indicative of cell polarization and morphogenesis. Moreover, we identified multiple myelin Schwann cell subtypes, two of which were enriched with mesenchymal differentiation genes or collagen-specific genes rather than putative myelin producing pathways. Across both the spontaneous partial denervation and surgical nerve crush injury models, SPP1 signaling emerged as an important mediator of interactions between myelin-producing Schwann cells, phagocytic cells, and tSCs, consistently driving Schwann cell proliferation associated with successful reinnervation. These findings underscore the universal importance of SPP1 signaling in NMJ dynamics and highlight it as a potential target for enhancing NMJ remodeling and preservation of muscle fibers. Our study offers novel insights into the biology of muscle-resident glial cells, paving the way for innovative therapeutic strategies in denervating conditions like amyotrophic lateral sclerosis (ALS) and age-associated skeletal muscle wasting.

## Results

### scRNA-Seq captures muscle-resident Schwann cells across healthy and denervated muscles

Activation of Schwann cell regenerative responses are consistently triggered by axonal injury, but the extent and persistence of the injury can lead to a spectrum of cellular processes within the regenerative timeframe. Utilizing a sciatic nerve crush injury model in Schwann cell reporter mice (S100GFP-tg), we induced complete muscle denervation, characterized by a total loss of isometric force production in gastrocnemius muscles upon nerve stimulation at 7 days post-injury (7 dpi) (Fig 1A). In uninjured 2-month-old *Sod1*^-/-^ mice, the force generated during nerve stimulation was approximately 33% lower than that of *Sod1^wt/wt^* controls while the force generated in response to direct muscle stimulation did not differ between genotypes (Fig. 1B). This finding is indicative of partial denervation in *Sod1*^-/-^ mice with the presence of muscle fibers that were highly contractile but non-responsive to nerve stimulation. Consistent with published reports of elevated markers of muscle fiber denervation at 2 months of age in *Sod1*^-/-^ mice (*10*), the neurotransmission impairments we observed in *Sod1*^-/-^ mice (crossed with the Schwann cell reporter, S100GFP-tg*Sod1*^-/-^ mice) were specifically noted at 2 months of age. The denervation event observed in the muscles 2 month old *Sod1*^-/-^ mice appears to be transient in nature as evidenced by the lack of differences in force elicited by nerve and direct muscle stimulation at either 1 or 3 months (Fig 1C). In contrast, there was no functional evidence of denervation in *Sod1^wt/wt^*controls at any time point evaluated (Fig 1C). This temporal pattern of innervation loss and subsequent muscle fiber reinnervation in muscle from *Sod1*^-/-^ mice correlates with direct assessments of NMJ denervation observed histologically, where the percentage of denervated NMJs was significantly elevated at age 2 months (∼23%) and returned to baseline by age 3 months (Fig. 1D). Morphological assessments of NMJs in both denervation models and healthy innervated controls confirmed the presence of tSCs across all conditions (Fig. 1E). Despite the varying degrees of denervation between models, tSCs remained a constant feature at NMJs, suggesting underlying transcriptional commonalities among the groups that warranted further investigation.

**Fig. 1.**
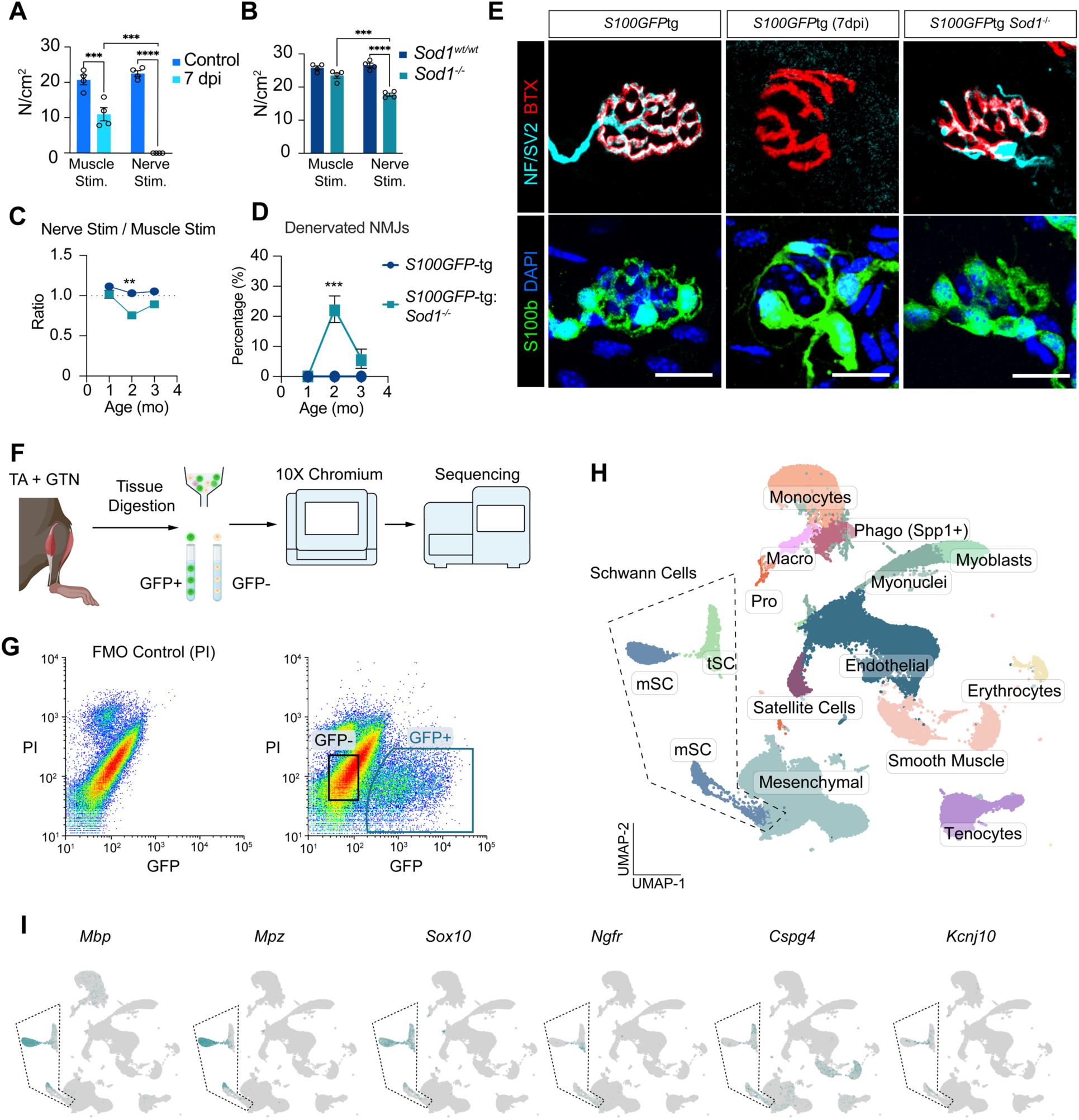
Characterization of muscle-resident Schwann cell subtypes across different denervation states via single-cell RNA sequencing (scRNA-Seq). (**A** and **B**) Maximum tetanic contraction forces (N/cm2) generated by nerve stimulation and direct muscle stimulation in WT mice 7 days post sciatic nerve crush injury (7dpi) (**A**) and in S100GFP-tg Sod1-/- mice (**B**), compared to age-matched WT controls (n = 4 per genotype). (**C**) Comparison of maximum isometric force ratios elicited by nerve versus direct muscle stimulation across 1-3 months in S100GFP-tg Sod1-/- (n = 3-5) and control mice (n = 3-5). (**D**) Percentage of denervated NMJs in gastrocnemius muscles of S100GFP-tg and S100GFP-tg Sod1-/- mice aged 1-3 months. (**E**) Representative staining of NMJs showing Schwann cells (S100B; green), nerve terminals (NF/SV2; cyan), acetylcholine receptors (AChR; red), and nuclei (DAPI; blue) in 2-month-old S100GFP-tg control and Sod1-/- mice, plus S100GFP-tg mice 7 days post-injury. (**F**) Experimental workflow: Bilateral gastrocnemius (GTN) and tibialis anterior (TA) muscles were harvested from 2-month-old S100GFP-tg, S100GFP-tg Sod1-/-, and S100GFP-tg mice 7 days post nerve injury, followed by FACS to isolate GFP+ and GFP-cells, then processed for scRNA-Seq using the 10X Chromium platform. (**G**) FACS plots showing gating strategies for GFP+ PI-single cells, with FMO (PI only) controls on the left. (**H**) UMAP plot visualizing 15 distinct cell clusters. (**I**) UMAP plots displaying transcript levels for myelin Schwann cell markers (Mbp, Mpz), general Schwann cell markers (Sox10), SC repair phenotype (Ngfr), and terminal Schwann cells (Cspg4, Kcnj10).

While the classification of Schwann cells has largely relied on histological examination and scRNA-Seq from nerve extracts (*11*), transcriptomic data on muscle-resident Schwann cells, especially tSCs, remains scant (*12–15*). A particular challenge faced by these studies is the rarity of tSCs and the extensive tissue requirement, limiting the cell numbers previously analyzed to as few as 100 individual cells. To overcome this limitation and gain a thorough understanding of the transcriptomic changes in muscle-resident Schwann cells, as well as other cells in the muscle microenvironment during NMJ remodeling, we used scRNA-Seq on pooled tibialis anterior (TA) and gastrocnemius (GTN) muscles from *S100GFP*-tg mice, *S100GFP*-tg *Sod1*^-/-^ mice, and *S100GFP*-tg mice 7 dpi, all at 2 months of age (Fig. 1F). After digesting the muscles, we sorted GFP+ PI- and GFP- PI- single cells using fluorescence-activated cell sorting (FACS) (Fig. 1G) and then subjected the cells to droplet-based scRNA-Seq. Our unbiased clustering approach using uniform manifold approximation and projection (UMAP) classified 54,273 cells [*S100GFP*-tg (n = 5,330); *S100GFP*-tg *Sod1*^-/-^ (n = 9,577); *S100GFP*-tg (7 dpi) (n = 39,366)] into 12 non-Schwann cell clusters and 2 Schwann cell clusters including myelin and terminal Schwann cell clusters (Fig. 1I). Both Schwann cell clusters (fig. S1A) displayed classic SC markers such as *S100b* and *Sox10* (Fig. 1J and fig. S1A) and no Schwann cell clusters were identified in GFP- cells. Notably, clusters of myelin Schwann cells were marked by the expression of myelin-associated genes like *Mbp* and *Mpz* (Fig. 1J, while the tSC cluster presented expression of recently reported non-myelin SC markers, *Cspg4* (*Ng2*) and *Kcnj10* (*Kir4.1*) (Fig. 1J) (*12*, *14*). Additionally, the tSC cluster prominently expressed *Ngfr* (p75NTR), a classical Schwann cell repair marker. *S100GFP*-tg and *S100GFP*-tg *Sod1*^-/-^ samples displayed a comparable total number of Schwann cells (1,649 and 2,101, respectively), however, relatively fewer Schwann cells (n = 369) were isolated from *S100GFP*-tg (7 dpi), despite having nearly 4-fold more total cells than the other groups. These muscles (7 dpi), however, were highly enriched for mesenchymal cells (+30%), monocytes (+10), and a *Spp1+/Cora1a+* cell cluster we termed as phagocytic (fig. S1b). Interestingly, we found many FACS GFP+ non-Schwann cell clusters that expressed *Gfp* mRNA but had little to no expression of *S100b* (fig. S2). These cells were abundant in *S100GFP*-tg (7 dpi) mice, and likely suggestive that Schwann cell de-differentiation was underway and prevalent at this injury timepoint, but this remains to be experimentally validated.

### Differential Schwann Cell Dynamics during NMJ remodeling

Re-clustering of Schwann cell transcriptomes revealed 5 subtypes including three myelin- SCs (mSC-A, mSC-B, mSC-C) and two distinct tSC subgroups, which we termed tSC-A and tSC- B (Fig. 2A-C). Notably, there was a significant increase in the tSC-B subgroup under both denervation conditions, representing ∼20% of the SCs in 2-month-old S100GFP-tg*Sod1*^-/-^ mice and ∼ 50% in the S100GFP-tg 7dpi mice but only 4% in the S100GFP-tg controls. Also common to both denervation models was a dramatic reduction in the proportion of myelin SC-A, representing 63% of the SCs in S100GFP-tg, but only 31% in S100GFP-tg*Sod1*^-/-^, and not detected in S100GFP-tg 7dpi mice as was also true for tSC-A. Additionally, the mSC-B and mSC-C clusters were more prevalent in the denervation models than in controls.

**Fig. 2.**
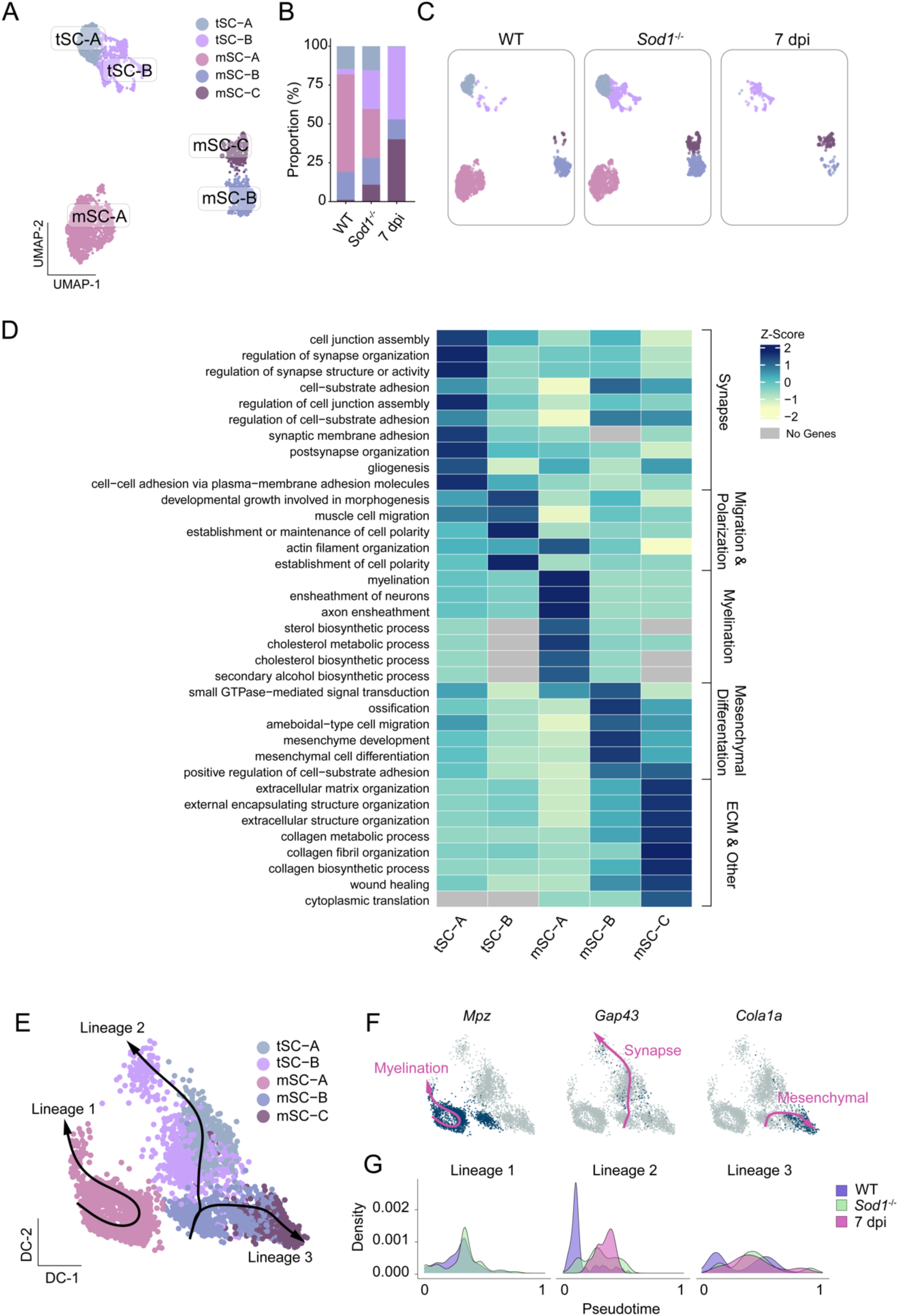
Dynamics of muscle-resident Schwann cell subtypes in healthy and remodeling neuromuscular fibers. (**A**) UMAP visualization of re-clustered muscle-resident Schwann cell clusters, identifying two terminal Schwann cell subclusters (tSC-A and tSC-B) and three myelin Schwann cell subclusters (mSC-A, mSC-B, mSC-C). (**B**) Proportions (%) of each Schwann cell subcluster, with corresponding UMAP plots (**C**) across conditions in WT, Sod1-/-, and 7 dpi mice. (**D**) Heatmap illustrating gene ontology (GO) pathway analysis results, with enriched biological processes for each Schwann cell cluster presented as z-scores of normalized -log(p-value) for each GO term. (**E**) Diffusion map showing Schwann cell subclusters with three trajectory lineages from slingshot analysis superimposed. (**F**) UMAP feature plots indicating expression patterns of key genes (*Mpz*, *Gap43*, *Cola1a*), highlighting their distribution within lineage 1, 2, and 3, respectively. (**G**) Variation in cell density over pseudotime for each lineage in WT, Sod1-/-, and 7 dpi mice, reflecting differential engagement in nerve repair processes.

Pathway analysis highlighted that the tSC-A cluster was associated with synapse organization and structural pathways (Fig. 2D, fig. S3A) while tSC-B showed enrichment for biological processes tied to migration, cell polarization, and morphogenesis (Fig. 2D, fig. S3B). The mSC-A cluster primarily engaged classical myelin production pathways, including axon ensheathment, myelination, and cholesterol metabolism (Fig. 2D, fig. S3C). Interestingly, our data suggest that mSC-B is characterized by gene expression programs defined by mesenchymal differentiation Fig. 2D, fig. S3D), consistent with findings from studies of peripheral nerve injury in which Schwann cells are reported to express epithelial-mesenchymal transition (EMT) markers and respond to TGFβ signaling, a well-known inducer of EMT genes (*16*, *17*). Finally, mSC-C was linked with extracellular matrix (ECM) organization and collagen production (Fig. 2D, fig. S3E).

To better understand the dynamic response of SC subpopulations during NMJ remodeling, we performed Slingshot trajectory analysis (*18*) on a diffusion map-reduced space of muscle-resident SCs (Fig. 2E). This analysis revealed three primary lineages. Lineage 1, which originates in the myelin Schwann cell cluster (mSC-A), demonstrated enrichment for myelin-associated genes such as *Mpz* (fig. S3C), underscoring its involvement in myelination and axonal support (Fig. 2E, F). The other two lineages arise from the mSC-B cluster and coursed through non-myelinating tSC subpopulations. Lineage 2 was characterized by an enrichment for synaptic-related genes, such as *Gap43* (Fig. 2 E, fig. S3A), highlighting its role in synaptic maintenance and regeneration. Meanwhile, Lineage 3 extended toward the mSC-C cluster, which showed enrichment for extracellular matrix (ECM) and mesenchymal differentiation genes, including COL1A1, suggesting its involvement in structural remodeling and repair (Fig. 2E, F, fig. S3E).

Our analysis across different experimental groups revealed a complete absence of myelination-associated Lineage 1 Schwann cells in WT (*S100GFP*-tg) mice 7 dpi. This finding aligns with established knowledge that acute nerve injury not only suppresses myelination-specific pathways but also leads to the expected reduction of Schwann cells contributing to these pathways, while concurrently activating myelin clearance, demonstrating a typical immediate response to nerve damage (*19*) (Fig. 2G). Conversely, control non-injured *S100GFP*-tg mice progressed more rapidly along the myelination trajectory, indicating robust myelination activity from the onset of the predicted trajectory. *Sod1*^-/-^ derived muscle-resident SCs, representing a partial denervation model, displayed a moderated progression in this lineage. Lineage 2, associated with synaptic maintenance and regeneration, revealed increasing cell density with pseudotime, beginning with the *S100GFP*-tg controls and progressively increasing in the *S100GFP*-tg*Sod1*^-/-^ and 7 dpi SCs. The pattern of increasing cell density in Lineage 2 is consistent with increased demands for synaptic remodeling in response to both partial and complete denervation through the remodeling trajectory. Lastly, Lineage 3, linked to ECM remodeling and mesenchymal differentiation, showed pronounced enrichment in the 7 dpi SCs, indicative of a shift towards repair functions following acute denervation. Both *S100GFP*-tg and *Sod1*^-/-^ SCs exhibited phases of activation within this lineage, indicating ongoing remodeling activities during neuromuscular adaptation and repair (Fig. 2G).

### The remodeling of neuromuscular junctions is associated with increased numbers of tSCs and larger synaptic areas

Given the substantial capture of muscle-resident Schwann cells in 2-month-old *Sod1*^-/-^ mice compared to the 7 dpi mice, and the recognition of this period as a crucial phase for Schwann cell-mediated repair against denervation, we undertook a focused examination of cellular and morphological changes during the denervation-reinnervation cycle in *Sod1*^-/-^ mice. We performed detailed imaging analyses on fixed fiber bundles from *S100GFP*-tg*Sod1*^-/-^ and *S100GFP*-tg control mice at 1, 2, and 3 months of age (Fig. 3A,B). Our comprehensive analysis defined NMJ structure, accounting for 16 distinct features related to pre- and post-synaptic structures including tSC number and morphology allowing us to track the dynamics of NMJ remodeling and Schwann cell activity over a transient repair period. Detailed descriptions of each morphological feature and how it is derived are described in fig S4. In muscles from 2 month old mice, our findings revealed generalized smaller values for *S100GFP*-tg*Sod1*^-/-^ mice compared to fully innervated controls in nerve terminal perimeter and overlap of the nerve terminal with AChRs by 33% and 60%, respectively (Fig. 3C,D). Meanwhile, the area of AChRs was observed to be 30% larger in S100GFP-tg*Sod1*^-/-^ mice compared to controls, and the total tSC area and number of tSCs per NMJ were 60% and 3-fold greater, respectively, in the *S100GFP*-tg*Sod1*^-/-^ group (Fig. 3E). As a result of the large increase in tSC number, the terminal area per tSC was 38% smaller in *S100GFP*- tg*Sod1*^-/-^ mice. Given that both central and peripheral glia are reported to respond to neuronal injury by increasing cell number (*2*, *20*), increased tSC number and area at the NMJ are likely acting to protect or promote NMJ integrity during the early neuromuscular events linked to the *Sod1*^-/-^ NMJ remodeling phenotype.

**Fig. 3.**
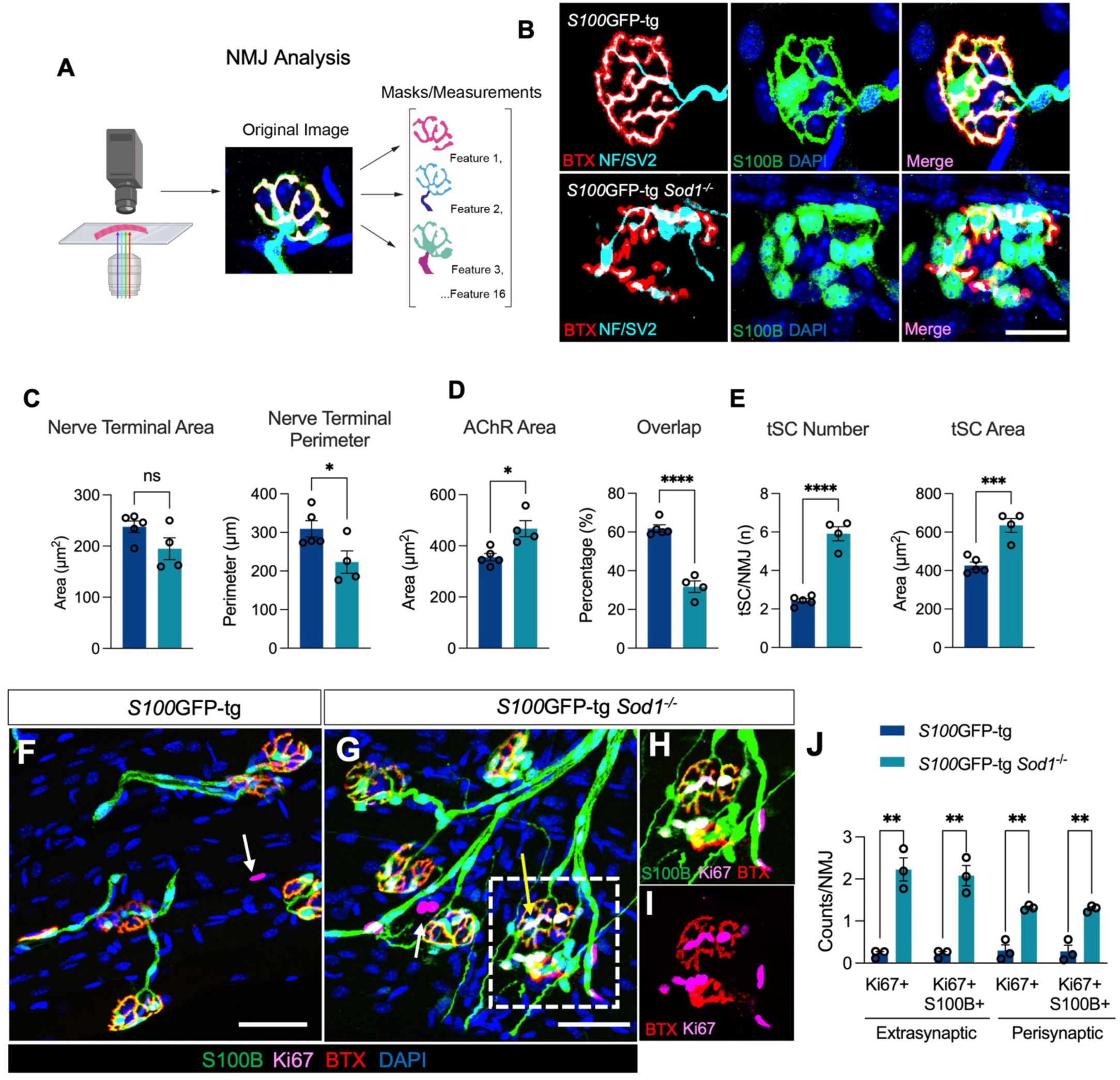
The remodeling of neuromuscular junctions (NMJs) is associated with greater tSC numbers, larger synaptic areas, and enhanced proliferation in *S100GFP*-tg *Sod1*^-/-^ mice. (**A**), Schematic of NMJ analysis – consisting of collecting NMJs images from muscle fiber bundles followed by the generation of feature masks and their measurements. Additional details on the generation of the masks are provided in fig S3 or 4 and Methods. (**B**), NMJs stained with S100B (Schwann cells; Green), NF/SV2 (Nerve; Cyan), α-bungarotoxin (AChR; Red), and DAPI (nucleus; Blue) in 2-month-old *S100GFP*-tg control and *S100GFP*-tg *Sod1*^-/-^ mice. (**C**-**E**) Quantification of nerve terminal area, nerve terminal perimeter, AChR area, percentage overlap between AChR area and nerve terminal area. and tSC number and tSC area. Muscle fiber imaging from S100GFP-tg (**F**) and S100GFP-tg Sod1-/- (**G**) mice, immunostained for S100B (green), Ki67 (magenta), α-bungarotoxin (red), and DAPI (blue). White arrows point to extrasynaptic nuclei positive for Ki67 but lacking endogenously expressed GFP and or S100B immunostaining, while the yellow arrow highlights a perisynaptic Ki67+GFP+ nucleus. (**H**) Enlarged view of two NMJs from the highlighted region in **G**, detailing the S100B, Ki67, and BTX stains. (**I**) The same NMJs from **H**, but focused on BTX and Ki67, revealing multiple Ki67+ nuclei in close proximity to the endplate. (**J**) Quantification of extraysnaptic and perisynaptic nuclei either singly labeled for Ki67 or double labeled for Ki67 and GFP. Values for all features across all NMJs analyzed are provided in fig S4. Open circles indicate average for each individual mouse of no fewer than 20 NMJs analyzed per muscle and bars represent means across animals ± SEM. *Denotes p < 0.05 vs *S100GFP*-tg *Sod1*^-/-^ by two tailed unpaired t-test. Scale bars for **B** equals 25 μm. Scale bars for **F** and **G** equals 50 μm.

Both the degree of overlap and the size of the synaptic area serve as indicators of the physical congruence of the pre- and post-synaptic components (*21*). These markers are robust histological indicators of nerve-to-muscle connections. In *S100GFP*-tg *Sod1*^-/-^ mice, the overlap between the nerve terminal and acetylcholine receptors (AChRs) was markedly decreased at 2 months compared to controls. However, by 3 months, this overlap in *S100GFP*-tg *Sod1*^-/-^ mice aligned with that of the *S100GFP*-tg controls (fig S5. Re-establishment of control levels of overlap could be explained by presynaptic responses, compensatory postsynaptic changes, or a combination. The present analysis revealed an expansion in synaptic area between 2 and 3 months in *S100*GFP-tg*Sod1*^-/-^ mice driven largely by an enlargement in nerve terminal area at the age of 3 months (fig S4). Collectively, the restoration at 3 months in the degree of overlap coupled with the reduction in denervated NMJs (characterized by NMJs with less than 10% overlap) (Fig 1D) strongly support reinnervation.

Based on our findings that NMJs in 2-month-old *Sod1*^-/-^ mice were characterized by higher numbers of tSCs than observed in other groups, we aimed to verify the presence of proliferation of *S100*GFP+ cells at the NMJs in these mice. To directly test this, we stained muscle bundles with an anti-Ki67 antibody (Fig. 3F-I). We found large numbers of Ki67+ nuclei in the fiber bundles from *Sod1*^-/-^ mice compared to age matched WT mice. Moreover, the majority of Ki67+ nuclei originated from cells that were also *S100*GFP+ positive. Ki67+ cells were observed both in perisynaptic (overlapping AChR staining) and extrasynaptic locations (within 50 microns of the nearest NMJ) (Fig 3J). These observations corroborate our earlier finding of a 3-fold increase in tSCs, in *Sod1*^-/-^ mice at two months of age when compared to wild-type controls (Fig. 3e).

Our functional and morphological data strongly support the existence of a key regenerative window centered around 2 months of age in *Sod1*^-/-^mice that is ideal for studying the cellular and molecular changes in muscle Schwann cells during NMJ remodeling and reinnervation. We also present compelling evidence that this remodeling process includes, at least in part, proliferation of Schwann cells located in the muscle at the NMJ and a promotion of increased synaptic area through nerve terminal growth. Thus, we next aimed to define key intercellular signals that regulate Schwann cell dynamics during the remodeling of the NMJ.

### Intercellular communication network analysis reveals a novel SPP1 signaling dynamic between myelin SCs and terminal SCs

Based on the unique location of tSCs at the NMJ, signaling to these cells is key to understanding the dynamics of this specialized environment. To explore how tSCs interact with other cellular components within the skeletal muscle niche during NMJ remodeling, we conducted an intercellular communication network analysis employing CellChat (*22*). Utilizing our scRNA sequencing data from tSCs, mSCs, mesenchymal progenitors, macrophages, and smooth muscle cells we identified shared secreted signaling pathways that target tSCs in all experimental groups: *S100*GFP-tg, *S100*GFP-tg *Sod1*^-/-^, *S100*GFP-tg (7 dpi), and a combined Denervation group encompassing both S100GFP-tg *Sod1*^-/-^ and S100GFP-tg (7 dpi) models (Fig. 4A). Transforming Growth Factor Beta (TGFβ) and Secreted Phosphoprotein 1 (SPP1) signaling emerged as the pathways with the highest normalized communication probabilities in the Denervation group, implying significant roles in NMJ remodeling in both partial and complete denervation conditions.

**Fig. 4.**
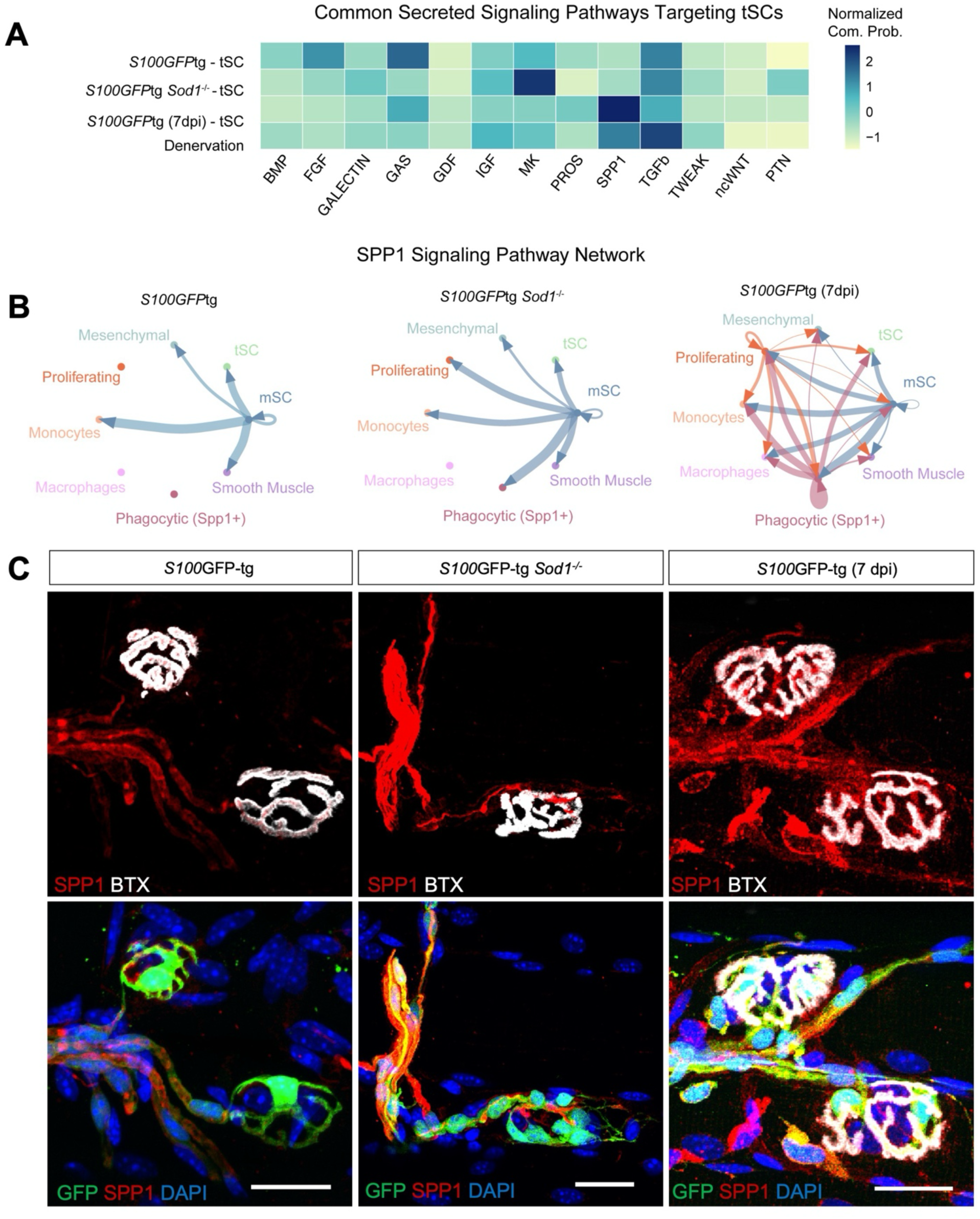
Intercellular communication infers an SPP1 signaling dynamic between mSC and tSC. (**A**) Heatmap showing the common significantly enriched secretion signaling pathways across WT, Sod1-/-, 7 dpi, and Denervation (Sod1-/- + 7dpi) targeting tSCs. Coloring of the heatmap is based on the normalized (*z*-score) communication probabilities. (**B**) Circle plots displaying the SPP1 signaling network across cell clusters for *S100GFP*-tg, *S100GFP*-tg *Sod1*^-/-^ mice and *S100GFP*-tg (7 dpi). The thickness of connecting lines represents the communication likelihood between paired cell clusters, with arrowheads demarcating communication directionality. (**C**) Representative immunofluorescent images of NMJs stained with SPP1 (Red), endogenous GFP (Green), BTX (White), and DAPI (Blue).

Our finding of marked induction of SPP1 signaling in our denervation groups is compelling in light of a recent study finding that SPP1 promotes Schwann cell proliferation and survival, and its expression is notably upregulated in myelin Schwann cells following human peripheral nerve injury (*23*). Exploring SPP1 signaling in our dataset, we identified a predominant expression of *Spp1* in mSCs (Fig. 4B). While SPP1 appears to communicate with several cell types in *S100*GFP- tg mice, a greater number of cell type receivers was inferred in *Sod1*^-/-^ mice (5 vs 7) (Fig. 4B), which included the addition of proliferating and Phagocytic (Spp1+) cells. Data from S100GFP-tg (7 dpi) mice revealed inferred SPP1 signaling originated from Phagocytic (Spp1+), myelin SCs, and Proliferating cell clusters, while the Phagocytic (Spp1+) cell cluster in *Sod1*^-/-^ mice was not predicted to communicate via SPP1 with other cells likely owing to the limited number of these cell identified in these mice. The SPP1 signaling was primarily predicted to act through *Cd44*, and various integrin dimer combinations with *Itgav*, *Itgb1*, *Itgb3*, *Itgb5* (fig. S5A).

To validate our CellChat findings, we performed targeted RT-qPCR to evaluate the expression levels of key genes within the SPP1 pathway in GTN muscles isolated from 2-month-old *S100GFP*-tg and *S100GFP*-tg *Sod1*^-/-^ mice (fig. S5B). We showed upregulation in the expression of *Tgfb1*, *Tgfbr2*, and *Spp1* in *Sod1*^-/-^ mice. While *Cd44* levels remained unchanged, we detected a borderline significant elevation in *Cd44v6* (p = 0.052), the specific receptor variant of CD44 known to bind SPP1. We next sought to confirm our scRNA-Seq and bioinformatic findings by pinpointing the protein localization of SPP1 in muscle fibers. Histological analysis using an Spp1 antibody on fixed muscle fiber bundles revealed pronounced localization of Spp1 associated with GFP+SCs near the NMJ, with intensified staining observed in *Sod1*^-/-^ mice compared to controls (Fig. 4C). In contrast to muscles from control and 2-month-old *Sod1*^-/-^ mice, in *S100GFP*-tg (7 dpi) SPP1 protein localization was observed in both GFP+ cells and GFP- negative cells. It seems likely that the GFP- cells represent the Spp1+ cell clusters in our scRNA-Seq dataset. Immunostaining also confirmed the localization of CD44 protein at the NMJ and within S100B+ Schwann cells (fig. S5C). Taken together, our findings illuminate a novel SPP1 cell signaling pathway involving myelin SCs, Phagocytic (SPP1+) cells and tSCs, which could play a role in enhancing the tSC proliferation and survival during muscle denervation important for successful NMJ remodeling and reinnervation.

### SPP1 gene expression is markedly increased in muscles following nerve injury

To explore whether an acute recoverable nerve injury alters the gene expression dynamics of SPP1 signaling in skeletal muscle, we performed sciatic nerve crush procedures on 2-month-old mice (Fig. 5A). Subsequently, we assessed transcript levels of denervation response pathways and genes involved in SPP1 signaling in GTN muscles in naïve non-injured controls, and 7-, 14-, and 28-days dpi. We observed a striking elevation in *Spp1* expression, and its receptors *Cd44* and *Itgav*, peaking at 7 dpi and subsequently reverting to baseline levels by 14 dpi (Fig 5A). This temporal trend closely parallels the known post nerve injury gene expression pattern of Chrna1(*9*), suggesting an inverse relationship with muscle innervation. In addition, Tgfb1 and its receptor Tgfbr2, proposed to be upstream of SPP1 signaling were also elevated at 7 dpi, with Tgfbr2 expression being reduced at 28 dpi compared to uninjured controls. Gene expression for NGFR and GDNF, which are associated with Schwann cell mediated nerve regeneration, were also highly expressed at 7 dpi, while the cell proliferation gene, CCND1, was elevated at 7 dpi and persisted at high levels out to 14 dpi. The pronounced *Spp1* expression during the initial nerve regeneration phase and its subsequent normalization is consistent with known muscle reinnervation milestones and implicates the potential involvement of SPP1 in muscle reinnervation.

**Fig. 5.**
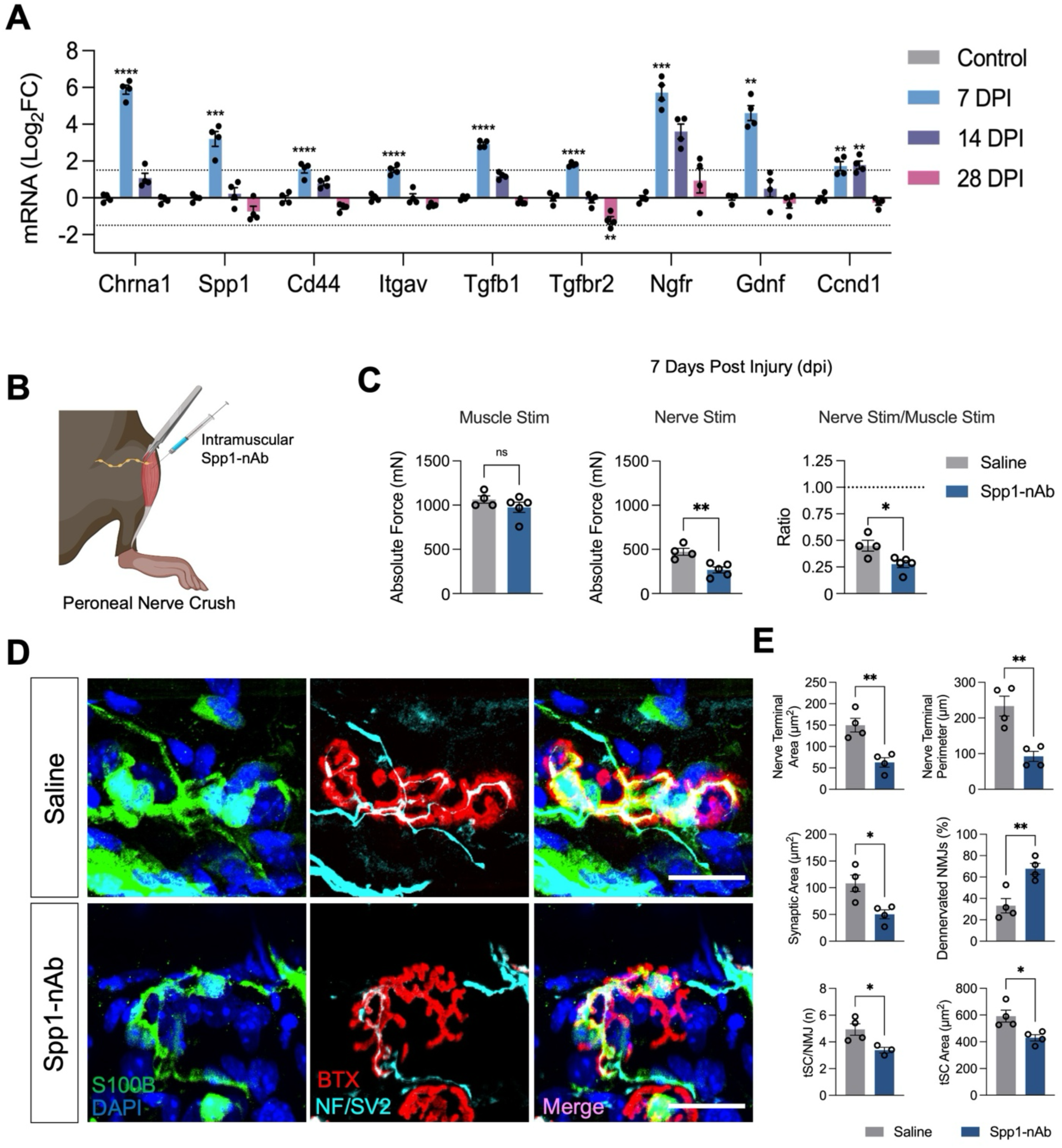
SPP1 signaling promotes tSC proliferation and muscle fiber reinnervation after nerve injury. (**A**) Following sciatic nerve crush injuries, gastrocnemius (GTN) muscles from C57BL/6 mice were collected at 0 (Control), 7, 14, and 28 days post-injury (**dpi**). mRNA expression levels of denervation markers (Chrna1), components of SPP1 signaling (*Spp1*, *Cd44*, *Itgav*, *Tgfb1*, *Tgfbr2*), genes linked to SC-mediated nerve regeneration (*Ngfr*, *Gdnf*), and cell proliferation (*Ccnd1*) at each time point are presented. (**B**) Peroneal nerve injuries were induced, and tibialis anterior (TA) muscles were intramuscularly injected with either Spp1-nAb or saline (control) at time of injury and every 2 days thereafter. (**C**) Data are shown for force (mN) evoked by direct muscle stimulation, with nerve stimulation, and ratio of force elicited by nerve and direct muscle stimulation at 7 **dpi** for saline- (gray) and Spp1-nAb-treated (blue) groups. (**D**) Representative immunofluorescent images of NMJs at 7 **dpi** stained with S100B (green), DAPI (Blue), BTX (Red), and NF/SV2 (Cyan). (**E**) Quantification of NMJ nerve terminal area and perimeter, synaptic area, percentage of denervated (>10% overlap) NMJs, tSC number and area. Open circles indicate values for individual mice and bars represent the mean across animals ± SEM. Scale bars represent 25 μm. *P ≤ 0.05, **P ≤ 0.01, ***P ≤ 0.001, ****P ≤ 0.0001 by two tailed unpaired t-test (Saline vs Spp1-nAb) or 1-way ANOVA (7, 14, 28 dpi vs Control) for multiple comparisons.

### Inhibition of muscle Spp1 after acute nerve injury results in reduced muscle reinnervation and fewer tSCs

To determine the role of Spp1 signaling in muscle reinnervation and its potential mediation of tSC responses post-nerve injury, we carried out peroneal nerve crush injuries on control *S100GFP*-tg mice and administered intramuscular injections of either an SPP1 neutralizing antibody (Spp1-nAb) or saline (Fig 5B). To expedite the evaluation of the onset of muscle reinnervation and reduce the number of muscle injections, nerve injuries were strategically performed near the nerve entry point of the TA muscle, with muscles receiving injections at the time of injury and every two days thereafter. Evaluations of recovery of functional neurotransmission as assessed by the ability of nerve stimulation to activate muscle contractions and tSC characteristics were conducted at 7 dpi. Evoked force measurements by direct muscle stimulation showed no difference between the saline- and Spp1-nAb-treated groups; however, nerve evoked muscle forces were 43% lower for muscles of mice treated with Spp1-nAb (Fig 5C). Furthermore, nerve-to-muscle force ratios in Spp1-nAb administered mice were 38% lower than for muscles of saline-treated mice. Collectively, these findings suggest that functional reinnervation post-injury was either impaired or delayed by neutralizing intramuscular Spp1 signaling.

We concurrently evaluated the effects of intramuscular SPP1 neutralization on tSCs in fixed muscle fiber bundles following injury histologically (Fig. 5E). Examination of tSC characteristics revealed a diminished number of tSCs per NMJ and a smaller tSC area for muscles that had SPP1 signaling inhibited (Fig 5E). In addition, mice treated with Spp1-nAb displayed a pronounced decrease in the nerve terminal area, perimeter, and synaptic area at 7 dpi in comparison to the saline-treated controls (Fig. 5E). Furthermore, the Spp1-nAb-treated group exhibited a significantly higher proportion of denervated muscle fibers relative to the saline-treated group (67% Spp1-nAb vs 33% saline), and NMJ synaptic area was also lower in Spp1-nAb mice. Taken together, this evidence, coupled with our functional analyses, underscores the role of Spp1 in muscle reinnervation and its involvement in augmenting the tSC number at the NMJ.

## Discussion

To our knowledge, the current study boasts the most detailed scRNA-Seq repository to date of muscle-resident Schwann cells, encompassing tSCs, across both healthy and denervated muscles. One prior study isolated tSCs using a dual mouse reporter that expressed both NG2- dsRed and S100GFP and performed bulk-RNA Seq (*12*, *24*), but bulk sequencing doesn’t allow for the distinction between different tSC subtypes or illuminate cell specific responses to NMJ remodeling. The results presented elucidate a previously unrecognized reactive population of tSCs that exhibits a transcriptional signature consistent with promoting synapse formation, enhancing cell adhesion, and organizing the extracellular matrix (ECM).

Our data also demonstrate tSCs are reactive, marked by increased proliferation underpinned by an intricate SPP1 signaling mechanism, bridging myelin Schwann cells and tSCs. Our observations of marked elevations in both SPP1 gene expression and protein immunofluorescence in the acute phase post muscle denervation is consistent with prior reports of induction of SPP1 in samples from nerve injury patients (*23*), and we show for the first time that SPP1 expression has functional relevance for the efficacy and/or efficiency of the response of tSCs to effect NMJ reinnervation. This significant conclusion is supported by our observations of increased tSC numbers following peroneal nerve injury that were notably decreased following intramuscular administration of an Spp1-nAb resulting in a diminished synaptic contact area, a pronounced percentage of denervated NMJs, and impaired recovery of functional innervation. Collectively, our findings suggest a pivotal role of SPP1 signaling, facilitated by myelin Schwann cells, in regulating tSC proliferation and reinnervation of denervated NMJs.

While upregulation of SPP1 in muscles of patients suffering from Duchenne Muscular Dystrophy (DMD)(*25*) and in mice following exercise (*26*) has been reported, our study uncovers a novel cellular origin contributing to this signaling — originating from muscle-resident myelin Schwann cells, rather than or perhaps in addition to macrophages targeting Fibro/adipogenic progenitors (FAPs)(*26–28*). The strong induction of SPP1 signaling pathway transcripts we observed concurrent with intense SPP1 immunofluorescence in skeletal muscle after nerve injury suggests SPP1 may have multiple functions to mediate muscle reinnervation. Integral to muscle reinnervation is Wallerian degeneration, a process governed by dynamic interactions among immune cells, FAPs, and Schwann cells (*29*). Thus, it is reasonable to hypothesize that SPP1 engages with multiple cell types in response to severe nerve trauma, such as nerve crush or nerve transection, to facilitate myelin clearance, ECM remodeling, and axon regeneration. In contrast, our observations, based on both imaging for protein and scRNA-Seq, SPP1 signaling was primarily inferred and observed from mSCs stemming in muscles of *Sod1*^-/-^ and was further elevated with complete denervation, with an additional signaling origin from SPP1+ Phagocytic cells. Furthermore, our findings of SPP1+ phagocytic cells, particularly following complete denervation, alongside the complete absence of classical myelin-producing Schwann cells, suggest a potential transition of Schwann cells to a phagocytic phenotype during denervation. Consequently, the SPP1+ cells observed at 7 dpi may originate from both Schwann cells and myeloid cells. This concept would integrate prevailing views in the field, which have seen a range of contradictory findings, of whether myelin clearance is primarily driven by SCs or through hematogenous macrophages (*30*). However, there is clear evidence that SC mediated phagocytosis is detected during the acute phase post-injury with hematogenous macrophages entering the injury site later (*31*, *32*) . Given the recent advancements in genetic mouse models and genomic technologies, this hypothesis warrants further investigation to clarify the roles and origins of these SPP1+ cells in the context of varying types of muscle denervation.

SPP1 signaling between glial cells is further supported by recent investigations showing elevated levels of SPP1 in nerve samples post-injury, which positively correlated with Schwann cell proliferation and survival (*23*). Furthermore, SPP1 has neuroprotective roles in the visual system and is expressed by reactive astrocytes to promote retinal ganglion cell survival after traumatic optic nerve damage (*33*). Beyond this regenerative aspect, SPP1 is implicated in pathological contexts, including tumor growth and metastasis, highlighting its versatility and likely the complexity of its regulation in promoting cell survival and growth in various contexts and cell types (*34*).

The synaptic regenerative signature we identified in tSCs during the response to denervation is distinctly characterized by the expression of *Gap43* and *Ntng1*. GAP43 is noted for its concentrated presence within regenerating growth cones of neurons and plays an indispensable role in axon guidance (*35*). The expression of GAP43 is not exclusive to neurons but is also found in reactive astrocytes and in tSCs after denervation, correlating with the elongation of tSC processes(*36*, *37*). Beyond axonal guidance, GAP43 is also involved in the transfer of mitochondria between astrocytes and glioblastoma cells (*38*). Given that mitochondria are abundant within growth cones to meet energy demands (*39*), one might speculate that *Gap43*+ tSCs promote axon regeneration through mitochondrial transfer, thereby contributing to metabolic support. This proposition opens new areas for exploring unconventional pathways of axon regeneration and delineating the multifaceted roles of GAP43 in neural repair mechanisms. Netrins are also important for axon guidance and netrin-1 has been shown to promote SC migration and proliferation (*40*, *41*). Recently, Netrin-G1was shown to facilitate signaling interactions between tSCs and sensory neurons, crucial for organogenesis in both hairy and non-hairy skin (*42*). The expression of *Ntn1g* in muscle-resident tSCs could function to support reinnervation of the endplate after denervation. Collectively, the expression of synaptic related genes in tSCs highlights the potential for new therapeutic targets to promote NMJ repair.

Cellular proliferation is universally recognized as a fundamental mechanism in various tissue regeneration processes. Although the proliferation dynamics of myelin Schwann cells have been extensively studied, with nerve injuries recognized as their primary trigger, the precise roles that these cells play during regeneration have not been established. A pertinent example is the marked 3 to 4-fold elevation in Schwann cell numbers observed within a week of sciatic nerve injury (*43*). While inhibition of Cyclin D1 mediated Schwann cell proliferation was ineffective in preventing nerve regeneration in the distal sciatic nerve stump (*44*), that study did not examine muscle-resident Schwann cells nor perform any functional assessment of NMJ transmission. tSCs not only proliferate after nerve injury but also extend cytoplasmic processes, promoting axonal sprouting, although the cellular signals remain under studied (*2*, *45*). Further highlighting the importance of tSCs for nerve regeneration, a recent study employed an anti-GD3 antibody that targets the GD3 ganglioside on tSCs, triggering complement-mediated lysis and resulting in targeted cell elimination after a peroneal nerve injury. This intervention led to a reduction in tSC numbers, decreased muscle reinnervation, and impaired functional recovery (*3*). Our study significantly expands on these previous studies by providing novel understanding of the role of SPP1-signaling acting on Schwann cells near the NMJ and its importance for the regeneration of NMJs.

Our research further highlights the *Sod1*^-/-^ mice as a useful model for understanding NMJ repair processes and tSC responses during the remodeling of innervation. Unlike nerve crush injuries, which denervate every muscle fiber leading to a lack of NMJ structural diversity, young *Sod1*^-/-^ mice recapitulate what we postulate to be a more faithful representation of heterogeneous NMJ remodeling in the context of normal aging and/or degenerative neuromuscular diseases. In support of this contention are our observations during the remodeling phase, of key characteristics reminiscent of NMJ regeneration, such as increased AChR area, tSC proliferation (*2*), polyinnervation (*46*), and axonal blebbing (*47*). Recognizing the potency of this model for investigating NMJ diversity and the response of muscle-resident Schwann cells to denervation informed our decision to perform scRNA-Seq, emphasizing GFP+ Schwann cells, which identified two distinct subtypes each for myelin Schwann cells and tSCs in muscles from 2-month-old KO mice and also 7 days post-nerve injury. A particular tSC subtype demonstrated traits related to ECM organization and synapse promotion and was considerably enriched in the *Sod1*^-/-^ muscles at 2-months of age, supporting their important role in NMJ regeneration.

Overall, our research deepens the existing knowledge on the intricate cellular processes of NMJ regeneration orchestrated by Schwann cells. Specifically, we highlight the importance of SPP1 signaling and elucidate the distinct roles played by specialized subtypes of muscle-resident Schwann cells. These insights not only expand our understanding of neuromuscular biology but also present promising avenues for targeted therapeutic strategies. Such innovations hold the potential to revolutionize the treatment landscape for debilitating neuromuscular disorders, including ALS and sarcopenia.

## Methods

### Animal Models

All experimental procedures were approved by the University of Michigan Institutional Animal Care and Use Committee (PRO00010548) and conducted in accordance with National Institutes of Health guidelines. Animals were sacrificed via intraperitoneal injection of tribromoethanol, followed by administration of a pneumothorax.

All mice used for experimentation were bred at the University of Michigan with *Sod1^−/−^* and WT mice initially provided generously by Dr. Holly Van Remmen while she was at the University of Texas Health Science Center at San Antonio. Mice expressing enhanced Green Fluorescent Protein (GFP) driven by the S100B promoter to visualize Schwann cells (S100GFP-tg Kosmos mice)(*48*) were generously provided by Dr. Alison Snyder-Warwick at Washington University in St. Louis, MO. *S100GFP*-tg mice were crossed with *Sod1^−/−^* mice to form a first generation of mice that contained a knockout allele for Sod1 and wild type allele for the Sod1 gene and the S100GFP transgene, S100GFP-tg *Sod1^+/−^*. First generation mice were crossed to generate a second generation of *Sod1^−/−^*mice expressing GFP under the S100B promoter to be used for our experiments. To maintain a stable colony of *S100GFP*-tg *Sod1^−/−^* mice, male *S100GFP*-tg *Sod1^−/−^* and female S100GFP-tg *Sod1^+/−^* mice were consistently bred to produce litters with approximately 50% of the mice containing the experimental genotype, *S100GFP*-tg *Sod1^−/−^*. Control mice in this study contain the genotype, S100GFP-tg *Sod1*^+/+^, simply referred to as S100GFP-tg.

### *In situ* Force Testing

Mice were anesthetized with initial intraperitoneal injections of Avertin (tribromoethanol, 250 mg/kg) with supplemental injections to maintain an adequate level of anesthesia during all procedures. Gastrocnemius (GTN) or tibialis anterior (TA) muscle contractile properties were measured in situ, as described by Larkin et al. (2011)(*49*). In anesthetized mice, the whole GTN or TA muscle was isolated from surrounding muscle and connective tissue using great care not to damage the nerve and/or blood vessels during the dissection. A 4-0 silk suture was tied around the distal tendon and the tendon was severed. The animal was then placed on a temperature-controlled platform warmed to maintain body temperature at 37°C. The hindlimb was secured to the platform, and the tendon was tied to the lever arm of a servomotor (model 6650LR, Cambridge Technology). A continual drip of saline warmed to 37°C was administered to the muscles to maintain temperature. Muscles were activated either by stimulation of the tibial (GTN muscle) or common peroneal (TA muscle) nerve using a bipolar platinum wire electrode or by direct stimulation of the muscle via cuff electrodes wrapped around the proximal and distal ends of the muscle. Custom-designed software (LabVIEW 2018; National Instruments, Austin, TX) controlled electrical pulse properties and servomotor activity and recorded data from the force transducer. The voltage of 0.2 ms stimulus pulses was increased, and optimal muscle length (Lo) was subsequently adjusted, to give maximum twitch force. The Lo was measured with digital calipers. Muscles were held at Lo and subjected to 300 ms trains of pulses with increase stimulation frequencies maximum isometric force (Po) was achieved. Muscle fiber length (Lf) was calculated based on previously established Lf-to-Lo ratios of 0.45 for GTN muscles and 0.6 for TA muscles. The physiological cross-sectional area (CSA) of muscles was determined by dividing the mass of the muscle by the product of Lf and 1.06 g/cm3, the density of mammalian skeletal muscle (*50*). Po was normalized by the CSA to calculate specific Po (sPo), as a measure of intrinsic force generating capacity. Functional innervation was assessed by calculating the ratio of force production in response to nerve stimulation relative to the force elicited by direct muscle stimulation. A value of 1.0 indicates high fidelity of the nerve-muscle functional connections, while the extent to which the ratio is less than 1.0 provides an indication of the fraction of muscle fibers that are unable to respond to nerve stimulation.

### Immunofluorescent Staining and Imaging

Animals were sacrificed and the muscles were fixed in 4% paraformaldehyde (PFA) in PBS for 15 min and sucrose protected overnight with 20% sucrose, followed by embedding in optimal cutting temperature (OCT) compound (Tissue Tek) and stored at -80°C prior to use. For visualization and examination of neuromuscular junction (NMJ) structure, muscles were dissected using fine forceps to tease muscle fibers into layers consisting of a maximum of 10-20 muscle fibers. The samples were then treated with ice cold methanol for 30 seconds, washed 3 times in PBS for 5 min each, then blocked for 30 minutes at room temperature in 5% goat serum, PBS, triton X-100 (0.5%), and 2% bovine serum albumin (BSA). Fab fragments (10% in blocking solution) were used to block endogenous IgG before applying any mouse monoclonal primary antibodies. Muscle fiber bundles were then stained with antibodies and mounted on a slide, and coverslips were applied.

Primary antibodies used for this experiment were as follows: a combination of NF and SV2 (Developmental Studies Hybridoma Bank) for labeling α-motor axon terminals and axons, anti- S100 (Dako, Z0311) to enhance labeling of all SCs, anti-Ki67 (rat), anti-SPP1 (Abcam, ab8448), and anti-CD44 (Thermo Fisher Scientific, 50-143-16). Secondary antibodies consisted of Alex Fluors that were conjugated to the secondary antibodies (Life Technologies). α-BTX conjugated to Alexa Fluor 555 (Invitrogen, B35451) was used to label the acetylcholine receptors (AChR), and 4′,6-diamidino-2-phenylindole (DAPI) was used to label all nuclei.

A Nikon A1 high-sensitivity confocal laser scanning microscope with 40× oil immersion objective was used to capture 16-bit, 512×512-pixel frame size, Z-stack images with 1 µm interval. A minimum of 20-30 NMJs were acquired per animal.

### NMJ feature quantification

Fiji software was used for quantitative analysis of confocal micrographs. All analyses were performed on maximum intensity projections using guidelines described in detail in the NMJ-Morph image processing pipeline (*21*). The obtained images were preserved as nd2 Nikon files and subsequently imported into ImageJ software, with channels consolidated into a single tiff image prior to their separation for individual component analysis. Initial preparations involved the division and cropping of images to ensure each contained a singular NMJ, enabling individual analysis of each NMJ. The analysis was constrained to en-face NMJs only.

For each channel, a threshold was set using the thresholding tool to produce a binary mask that most closely resembled the real image of the NMJ component being analyzed. The binary mask was selected and measured to provide the area and perimeter of the selected channel (fig S3). The endplate mask was generated by applying the ‘Create background’ FIJI tool on the AChR mask, which completely fills in the AChR mask. Endplate area and diameter were also measured (fig S3A). This process was repeated for every NMJ image, for each of the NMJ components.

The ‘Overlap’ and ‘SC Coverage’ measurements (fig. S4 D,E) were derived in accordance to NMJ-Morph guidelines. Briefly, the following calculations were used:

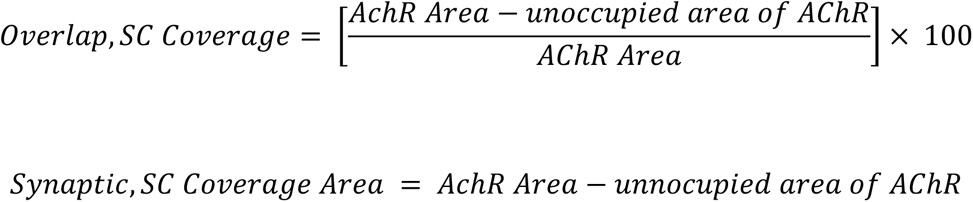

The unoccupied area denotes regions lacking colocalization with α-BTX. Following the generation of a binary mask for calculating the AChR, nerve teriminal, and tSC areas (Extended fig S3A-C), the masks were processed, inverted as needed, and merged to measure unoccupied areas of AChR. Beyond structural area measurements, attention was given to NMJs manifesting visible polyinnervation—characterized by the convergence of at least two axons onto a single NMJ (fig S3F). Furthermore, the presence of axon terminal blebs per NMJ was recorded (fig S3F).

### Quantification of Perisynaptic and Extrasynaptic Ki67+ Nuclei

Muscle fiber bundles from 2-month-old *S100GFP*-tg and *S100GFP*-tg *Sod1*^-/-^ were processed and stained with anti-Ki67, DAPI, α-BTX, and anti-S100B to delineate nuclei and cellular structures. Confocal imaging was performed using a Nikon A1 microscope, with images captured at 20X and 40X magnifications. The imaging focused on fields containing NMJs along with visualization of the surrounding areas. Classification of perisynaptic Ki67+ nuclei involved those being DAPI+ Ki67+ and overlapping with α-BTX stain. Perisynaptic Ki67+ GFP+ nuclei were identified as DAPI+ Ki67+ S100GFP+ and overlapping with an endplate (α-BTX stain). Extrasynaptic Ki67+ nuclei were those located within 50 μm from the nearest endplate. The quantification was normalized by the number of NMJs per image field to ensure comparability across different fields and conditions. A minimum of 50 NMJs were analyzed across each muscle.

### Single cell isolation via FACS

For tissue collection, mice were anesthetized with 3% isoflurane, then euthanized by cervical dislocation, bilateral pneumothorax, and removal of the heart. Hind limb muscles (TA and GTN) of control and experimental mice were quickly harvested using sterile surgical tools and placed in separate plastic petri dishes containing cold phosphate-buffered saline (PBS). Using surgical scissors, muscle tissues were minced and transferred into 50 mL conical tubes containing 20 mL of digest solution (2.5 U/mL Dispase II and 0.2% [∼1000 U/mL] Collagenase Type II, 1 mg/mL of hyaluronidase in Dulbecco’s modified Eagle medium [DMEM] per mouse). Samples were incubated on a rocker placed in a 37°C incubator for 60 min with manual pipetting the solution up and down to break up tissue every 30 min using an FBS-coated 10 mL serological pipette. Once the digestion was completed, 20 mL of F12 media containing 20% heat inactivated FBS was added into each sample to inactivate enzyme activity. The solution was then filtered through a 70 mm cell strainer into a new 50 mL conical tube and centrifuged again at 300g for 5 min. Live cells were sorted from the suspension via addition of 1 μg of propidium iodide (PI) stain into each experimental sample and all samples were filtered through 70 mm cell strainers before the FACS. Cell sorting was done using a BD FACS Aria III Cell Sorter (BD Biosciences, San Jose, CA) and BD Discovery S8 Cell Sorter (BD Biosciences, San Jose, CA). GFP+ PI- and GFP- PI- cells were sorted into 0.02% BSA PBS solution for immediate processing.

### scRNA sequencing

Freshly isolated single cells were sorted into staining solution, enumerated by hemocytometer, and re-suspended into PBS. Cells were loaded into the 10X Genomics chromium single cell controller for each sample and captured into nanoliter-scale gel bead-in-emulsions. cDNAs were prepared using the single cell protocol as per manufacturer’s instructions and sequenced on a NovaSeq instrument (Illumina) with 26 bases for read 1 and 159 bases for read 2.

### scRNA-Seq data processing and analysis

CellRanger 8.0.0 (10X Genomics) was used to process raw data using the GRCm39 reference with eGFP. The CellRanger workflow aligned sequencing reads to the GRCm38 transcriptome using the STAR aligner and exports count data^63^. The CellRanger `count` command was run with default parameters. Filtered feature barcode data were imported into Seurat (v5)(*51*). Ambient RNA was removed using Decontx (*52*) and only cells with greater than 200 genes and <10% mitochondrial reads were included in our analysis. Cell doublets were removed using DoubletFinder (*53*). Scaling, normalization, variable gene selection, dimensionality reduction, and clustering were performed with default settings using the Seurat. Batch integration across all samples was performed using Harmony (*54*), and cell clustering was performed using a resolution equal to 1. Cell types were assigned to each cluster using known marker genes. Pathway analysis was performed using enrichGO() function in clusterProfiler (*55*) with the default parameters and used gene markers that were upregulated (log2FC > 1) in the tSC clusters compared to all other muscle-resident Schwann. Upregulated genes were determined using the `FindMarkers()` function in Seurat. For each muscle-resident Schwann cell cluster, the top 10 biological processes (based on adjusted p-values) were aggregated into a single dataframe, retaining unique pathways only. Pathways were scaled as z-scores across the dataset to compare pathway enrichments across muscle-resident Schwann cells. A diffusion map was generated with the Destiny package (*56*), using selecting the first two diffusion components (DC) for detailed mapping, and Slingshot (*18*) was used for trajectory analysis with the omega parameter set to "`TRUE`" to allow for multiple lineage origins. Cell-cell interaction analysis was performed with CellChat, focusing on “Secreted Signaling” pathways to assess intercellular communication within the muscle-resident Schwann cell niche.

### Whole-tissue RNA extraction and RT-qPCR Analysis

GTN and TA muscle samples were homogenized in TRIzol reagent (Invitrogen, Thermo Fisher Scientific) using a bead mill. RNA was isolated by phenol/chloroform extraction followed by isopropanol precipitation according to the manufacturer’s protocol and RNA yield was determined using a NanoDrop Spectrophotometer (NanoDrop 2000c, Thermo Fisher Scientific). Genomic DNA was removed by incubation with DNase I (Ambion, Thermo Fisher Scientific, AM2222) followed by its heat inactivation. Total RNA (1 μg) was reverse-transcribed to cDNA using SuperScript III Master Mix, Random Hexamers, and dNTPs (Invitrogen, Thermo Fisher Scientific) and RT-qPCR performed on a CFX96 Real-Time PCR Detection System (Bio-Rad, 1855195) in triplicate 20 μL reactions of iTaq Universal SYBR Green Supermix (Bio-Rad, 1725124) with 1 μM forward and reverse primer. Relative mRNA expression was determined using the 2^-ΔΔCT^ method with *Gapdh* serving as a control for the samples.

### Nerve injuries and neutralization of intramuscular Spp1

Male C57BL/6J mice, aged between 10 - 16 weeks, were subjected to bilateral sciatic or peroneal nerve crush to induce muscle denervation in both legs. Fine # 5 forceps were employed to produce the nerve crush. Mice assigned to sciatic nerve injury protocols had tissues harvested at intervals of 0, 7, 14, or 28 days following the injury. For those undergoing peroneal nerve injury, a 10 μL intramuscular injection of SPP1 neutralizing antibody (4 μg, AF808, R&D Systems) was administered into the tibialis anterior (TA) muscles, utilizing a Hamilton syringe. This injection routine was repeated every two days post the initial injury. Control mice received saline injections concurrent with the day of injury and repeated every two days thereafter. Seven days post injury (**dpi**), all mice were anesthetized and TA muscles were assessed for contractile functionality and collected for NMJ imaging.

### Statistical analyses

Data are presented as the mean ± SEM. All statistical analyses were performed using GraphPad Prism 8 (GraphPad Software) and *R*. Between-group differences were tested by two-tailed unpaired students t-tests (2 groups) or by a one-way analysis of variance (ANOVA) followed by Dunnett’s multiple comparisons test. A two-way ANOVA followed by Tukey’s multiple comparison’s test was executed when two factors were involved. Differences were considered to be statistically significant at the *P* < 0.05 level (*p < 0.05, **p<0.01, ***p < 0.001, ns: not significant).

## Data availability

All raw and processed data are available on NIH GEO (GSE246865, GSE291457)

## Code availability

All source code will be available on GitHub: https://github.com/sdguzman/Muscle_SC

## Study Approval

All animal experiments were approved by the University of Michigan Institutional Animal Care and Use committee (IACUC)(PRO00008744).

## Acknowledgments

This work was supported by the National institutes of Health (NIH) under the awards R01(AG050676) (SVB), R01 (AG086251), P01 (AG051442) (SVB), P30 (AR069620)(SVB), T32 (AG000114)(SDG), and the American Physiological Society Porter Fellowship (SDG).

---

**fig. S1.**
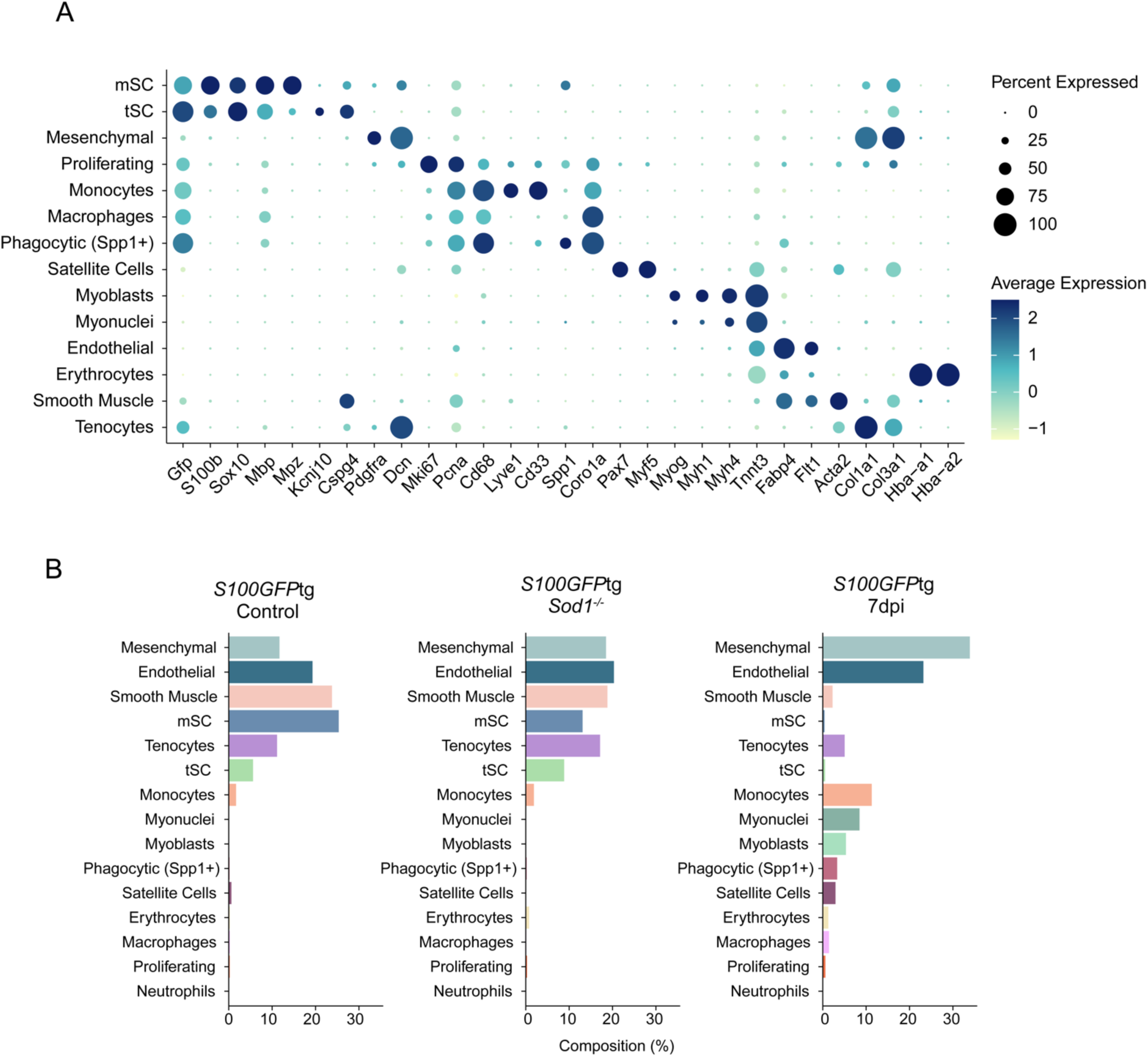
scRNA-Seq annotations and cell composition. (**A**) Dotplot showing the expression pattern of canonical markers, including *Gfp*, for each identified cell type cluster in the integrated dataset. (**B**) Bar plots showing the percentage (%) of each cell type cluster across all three groups.

**fig. S2.**
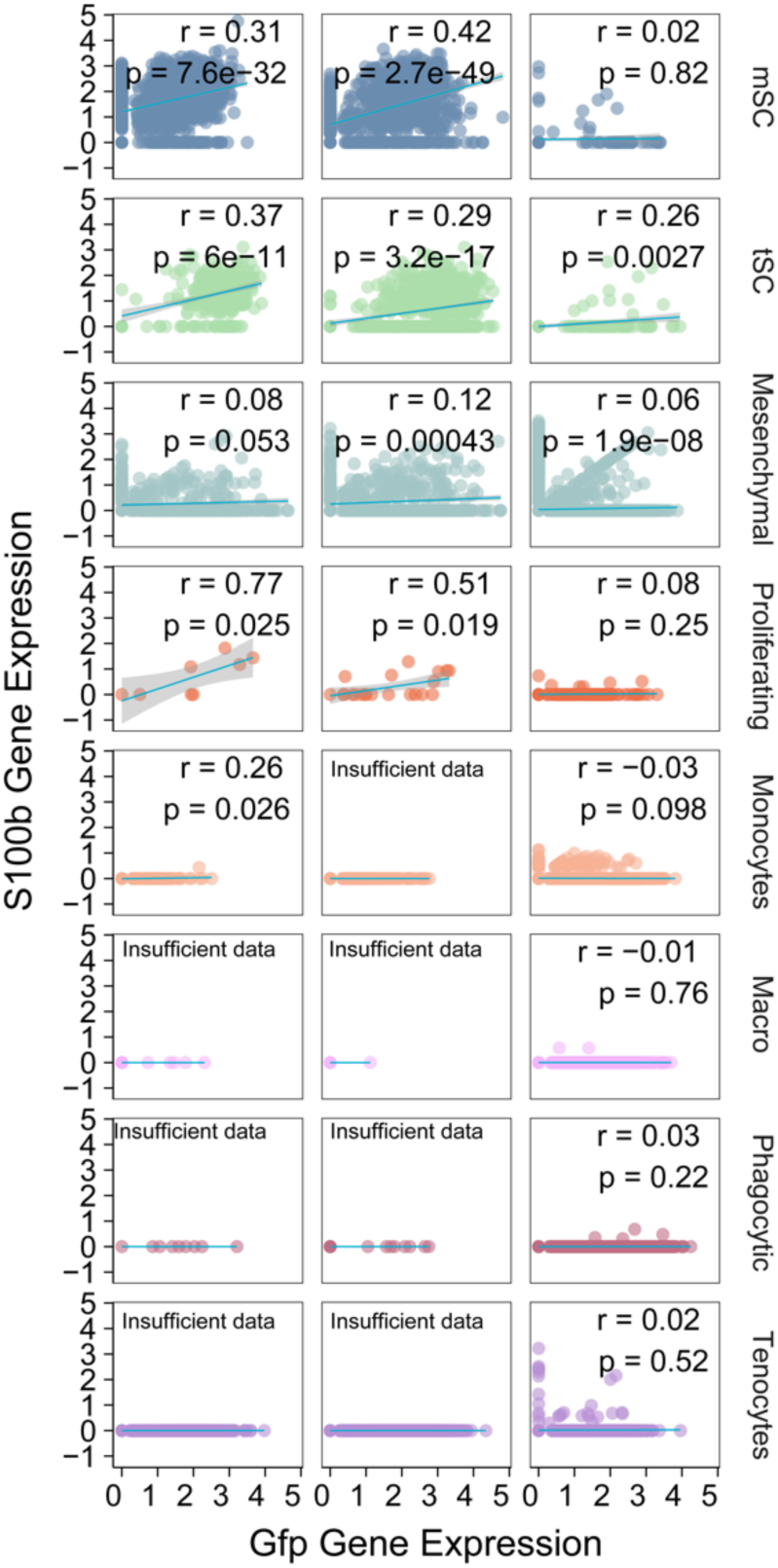
Correlation between Gfp and S100b expression across diverse cell types. Linear regression analysis of *Gfp* versus *S100b* expression in individual cells across various cell types within the integrated dataset that exhibited *Gfp* gene expression. Each subplot illustrates the regression line and includes the Pearson correlation coefficient, annotated where sufficient data points were available to perform the analysis.

**fig. S3.**
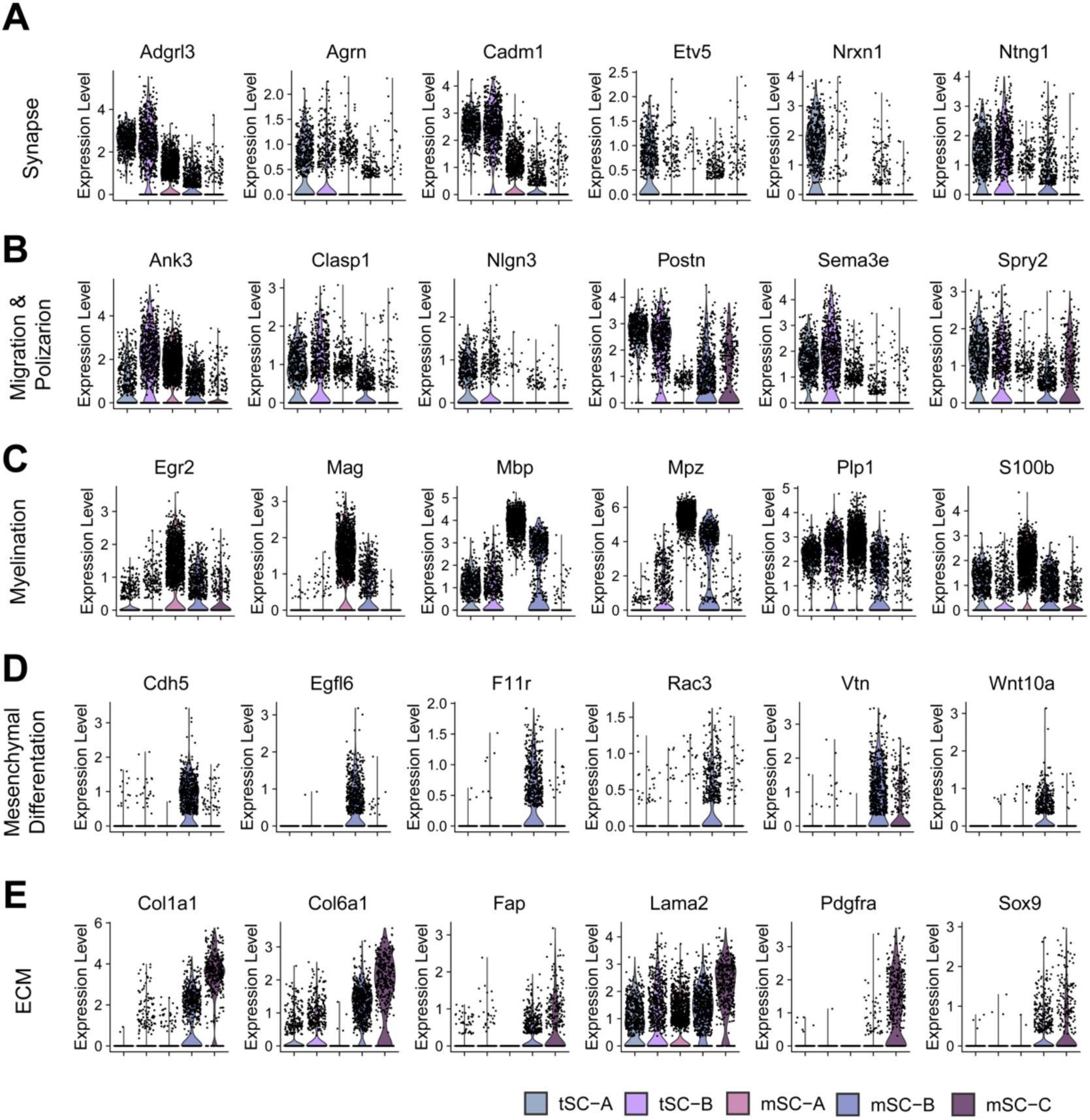
Differential expression of upregulated genes across pathway analysis categories in muscle-resident Schwann cells. Violin plots illustrate the expression patterns of enriched markers within the broad categories identified in Fig. 2—Synapse, Migration & Polarization, Myelination, Mesenchymal Differentiation, and ECM—across five muscle-resident Schwann cell subclusters.

**fig. S4.**
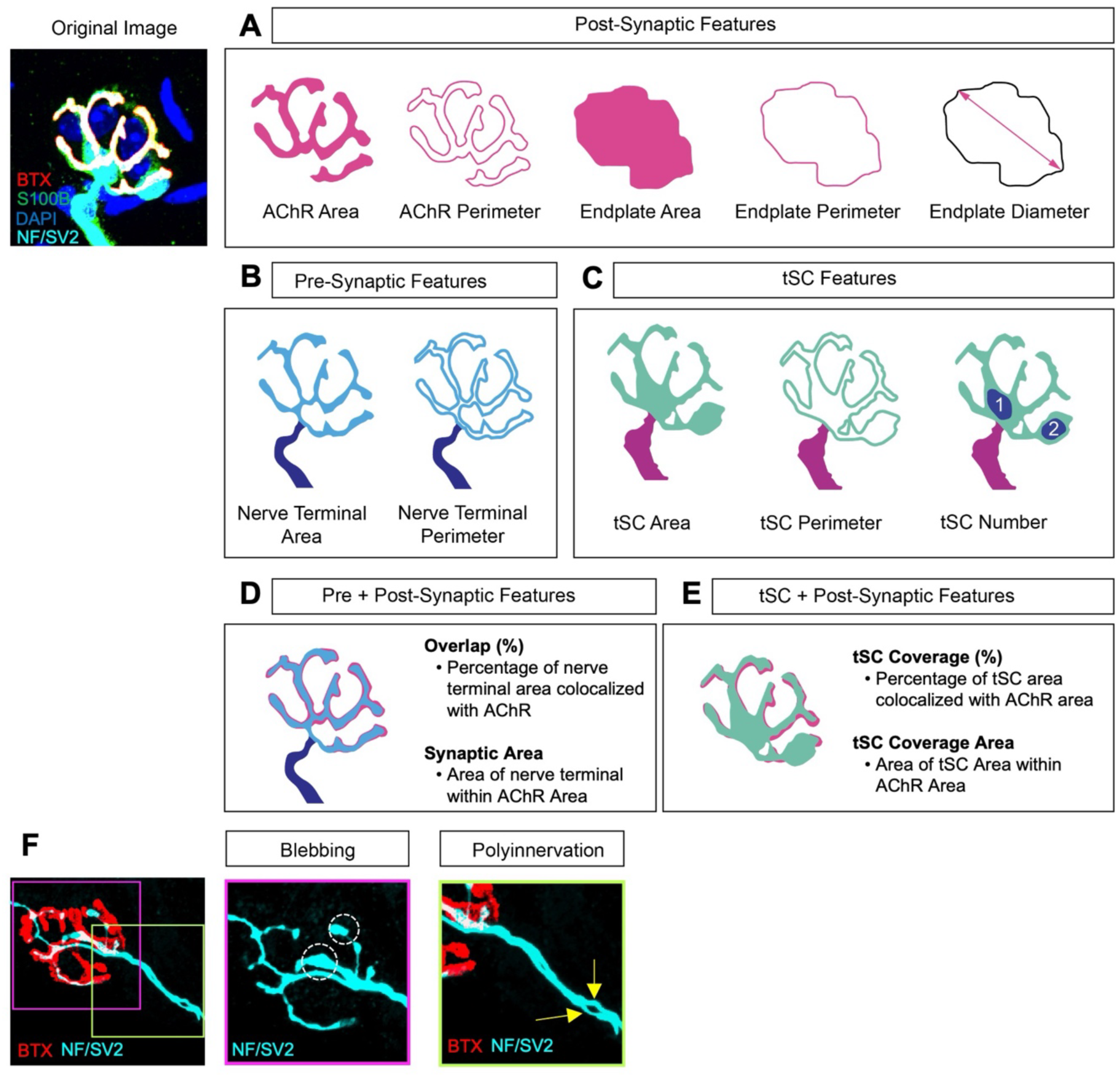
NMJ features analyzed. Binary masks were generated from the original image, and 16 en face NMJ morphological features were measured that represented the (**A**), post-synaptic (AChR and Endplate), (**B**), pre-synaptic (nerve terminal, axonal blebbing, polyinnervation), and (**C**), tSC structures. These pixel measurements are visualized by distinct colorations in each mask: red for post-synaptic features, light blue for pre-synaptic features, and green for tSC structures. (**D**), In addition, the percentage “Overlap” between axon terminal and AChR areas were measured. This overlap was further quantified in terms of synaptic area (μm^2^), denoting the area where the nerve terminal co-localizes with the AChR region. (**E**), Similarly, the “tSC coverage” was determined representing the tSC area that overlaps with the AChR region, and measured the "tSC Coverage Area," which is the expanse of tSC found within the AChR area. (**F**), The number of axon terminal blebs and and the presence of polyinnervation were also assessed for each NMJ.

**fig. S5.**
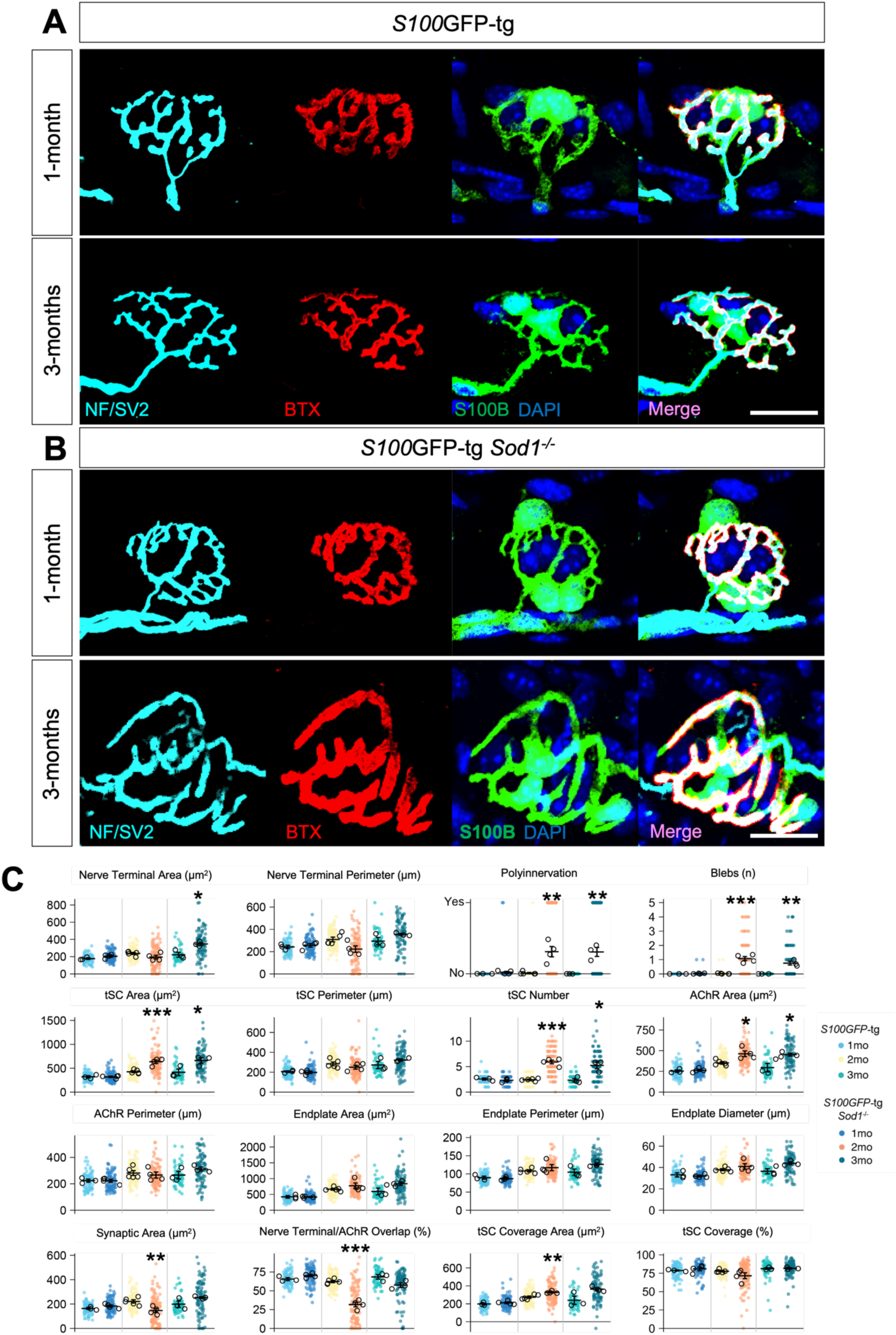
Analysis of NMJ in S100GFP-tg and S100GFP-tg Sod1-/- mice. (**A** and **B**) Images of representative NMJs from 1- and 3-month-old mice, stained for S100B (indicating Schwann cells in Green), NF/SV2 (Nerve in Cyan), bungarotoxin (AChR in Red), and DAPI (Nucleus in Blue). (**A**), Shows NMJs from S100GFP-tg mice, while (**B**) are NMJs from S100GFP-tg Sod1-/- mice. (**C**), Graphical representation of NMJ features in S100GFP-tg and S100GFP-tg Sod1-/- mice aged 1, 2, and 3 months. Each colored dot represents a single NMJ. Black circles denote the average for all NMJs in an individual animal. The horizontal bars show the overall mean for each group, while error bars illustrate the mean ± SEM. *Denotes p < 0.05 S100GFP-tg vs S100GFP-tg Sod1-/- by two tailed unpaired t-test. Scale bar equals 25 μm and images constrained to the same dimensions.

**fig S6.**
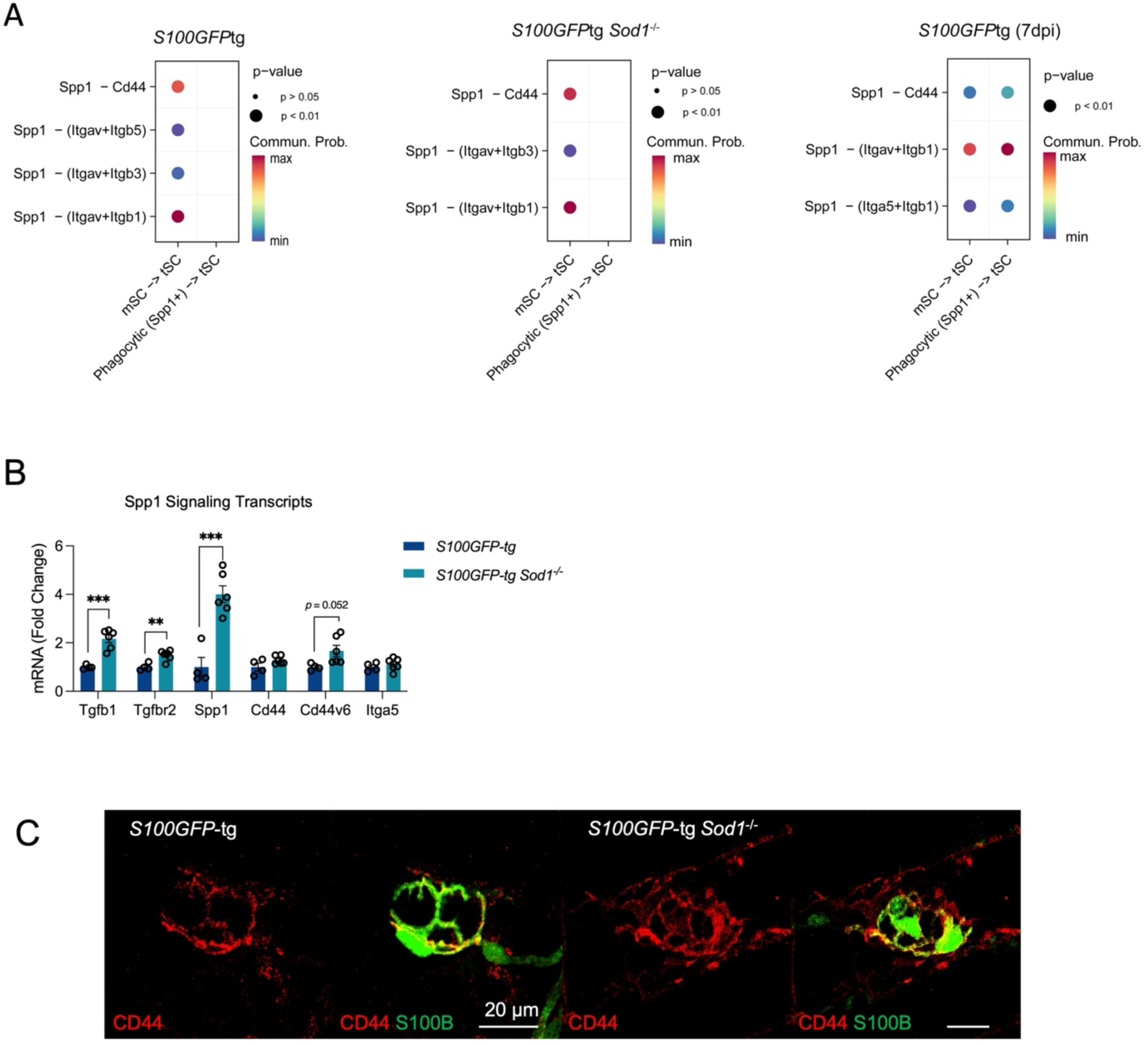
SPP1 signaling across different innervation states and confirmation of enhanced SPP1 gene expression and protein staining in *Sod1-/-* mice muscles. (**A**) Dotplot visualizing signaling of the Spp1 ligand to Cd44 and integrin receptors, denoting interactions between mSCs and tSCs in both *S100GFP*-tg and *S100GFP*-tg *Sod1*^-/-^ , *S100GFP*-tg (7 dpi) mice. (**B**) mRNA levels determined by qPCR for several presumed components of the Spp1 signaling pathway for muscles of *S100GFP*-tg (n = 4) and *S100GFP*-tg *Sod1*^-/-^ (n = 6) mice. (**C**) Representative immunofluorescent images of NMJs stained for CD44 (Red) and S100B (Green). Open circles indicate values for individual mice and bars represent the mean across animals ± SEM. Scale bars represent 20 μm. *p ≤ 0.05, **p ≤ 0.01, ***p ≤ 0.001, by two tailed unpaired t-test (S100GFP-tg vs S100GFP-tg Sod1-/-).

